# Image-based Morphological Profiling Reveals Signatures of Radiation Exposure

**DOI:** 10.1101/2025.11.27.691043

**Authors:** Olivia N Tiburzi, Laura L Dunphy, Sarah M Ton, Leah Talbott, Ryan J McQuillen, Alan W Hunt

**Affiliations:** Applied Biological Sciences Group, The Johns Hopkins University Applied Physics Laboratory, Laurel, MD, USA; Biological Sciences Group, The Johns Hopkins University Applied Physics Laboratory, Laurel, MD, USA; Applied Chemistry and Physics Group, The Johns Hopkins University Applied Physics Laboratory, Laurel, MD, USA

## Abstract

Exposure to ionizing radiation has the potential to induce significant health risks including radiation sickness and death. Here, the utility of image-based morphological profiling (IBMP) assays was investigated as a method to visualize signatures of radiation over time. Human-derived fibroblasts were used as the model system, and were exposed to varying doses of radiation. Cell Painting protocols were then applied to generate images for profiling. Quantitative analysis of images taken from fibroblasts exposed to 1 Gray of ionizing radiation revealed considerable morphological changes by 24 hours post-exposure, with some morphological signs of exposure emerging as early as 4 hours post-exposure. This work demonstrates proof-of-concept for the use of cell painting and IBMP to visualize signatures of radiation, paving the way for its use as a tool for assessing repeated exposures, screening of potential treatments, and establishing relevant timepoints for downstream orthogonal assays.

## INTRODUCTION

Ionizing radiation, a high-energy form of radiation capable of displacing electrons from atoms, poses a significant health hazard to humans^1^. While it has long been known that the health effects of radiation exposure are dose-dependent, and that acute exposures to high doses of ionizing radiation can induce radiation sickness, it has become apparent that even low-dose exposures (at or below 0.5 Gray (Gy)), can result in the onset of disease when repeated over time^2^. The link between this low-dose radiation (LDR) exposure and cancer incidence is well established, but more recent work has implicated LDR in the development of a variety of other adverse health conditions including immune system dysfunction, cardiovascular disease, neurological conditions, and cataracts^3^. This being said, the National Academies of Sciences, Engineering, and Medicine (NAAS) has suggested that this field of research is “limited and fragmented,” especially with regards to the exact molecular mechanisms by which LDR induces negative health effects^3^. Omics-based studies of LDR provide valuable snapshots of pathways impacted by radiation, but the expensive nature of these methods limits the number of conditions (*e.g.,* cell types, exposure doses, time points post exposure) that can be reasonably evaluated in a given experiment. The development of medium- to high-throughput pre-screening approaches to identify LDR exposures which reliably impact cell physiology would be valuable to inform both (i) the collection of orthogonal measurements (*e.g.,* transcriptomics, lipidomics), and (ii) avenues for the discovery of novel therapeutic interventions. Such pre-screening approaches could further be leveraged to temporally assess recovery following radiation exposure, as well as to study the impact of repeated LDR exposures.

Recently, image-based morphological profiling (IBMP) has shown promise as a strategy to rapidly repurpose and/or discover therapeutics for novel targets^4^. IBMP, also referred to as “phenomic analysis” or “morphological profiling,” is a low cost, high-throughput screening approach to compare and relate subtle changes in host cell phenotypes in response to extrinsic perturbations such as small molecules or soluble protein factors^4,5^. The input data for this approach is generated using Cell Painting, where cells exposed to a perturbant are fixed, stained with a panel of fluorescent dyes that label important subcellular structures, and imaged in high-throughput^6^. Following image acquisition, individual cells are identified (“segmented”) and over 1000 quantitative measurements are made per cell (*e.g.,* cell area, stain intensity, texture, and granularity). This process converts each image into hundreds of numeric snapshots, or cellular “phenoprints”^7,8^. To reduce noise, individual cell phenoprints can be aggregated to acquire well-level phenoprints, which can then be analyzed with unsupervised and supervised machine learning (ML) approaches. Such analyses allow for the comparison of the effects of different perturbations (*e.g.,* different doses of radiation), while also tracking cellular recovery over time, or in response to administration of countermeasure candidates.

Due to these advantages, phenomics-based approaches have already shown utility at multiple stages of drug discovery, including the identification of both novel small molecule therapeutics and chemical hazards in mammalian cell models^4,9–14^. Phenomics has also been shown to enable clustering of small molecule therapeutics and soluble immune stimuli by mechanism of action; however, to our knowledge, this method has not yet been used to examine the phenotypic signatures that arise from radiation exposure^9^. Here, human-derived fibroblasts were irradiated and an in-house phenomics pipeline was used to test the ability to identify exposures of varying doses at varying post-exposure time points. It was shown that this technique can reproducibly identify irradiated from unexposed cells, even at relatively low doses. This strategy may enable the use of IBMP as a pre-screening assay to prioritize conditions for follow-up with low-throughput, expensive mechanistic assays. Furthermore, it could additionally be adapted to assess the impacts of repeated LDR exposures, as well as to screen for treatments against radiation-based illnesses.

## RESULTS

### A system for the controlled irradiation of human-derived fibroblasts

An in-house pulsed radio frequency electron linear accelerator was optimized to reliably administer targeted doses of bremsstrahlung radiation to human-derived fibroblasts in 96-well microplates. Two replicate radiation experiments were performed, each testing two target doses (1 Gy, 3 Gy). For each exposure, three plates were stacked, irradiated together, and then placed back in the incubator for various periods of time (4, 24, 48 hours). During the exposure, one half of plates were shielded by lead to serve as the unexposed control samples. This methodology resulted in the reliable meeting of target radiation doses (**Table 1**). Stacking of three plates at a time did not considerably impact target dose and shielded wells were found to consistently receive on the order of 1-2% of the target dose (**Table 1**). Together, this system limits experimental variability by dosing multiple plates at a time and by including unexposed and exposed cells on the same plates.

**Table 1.**
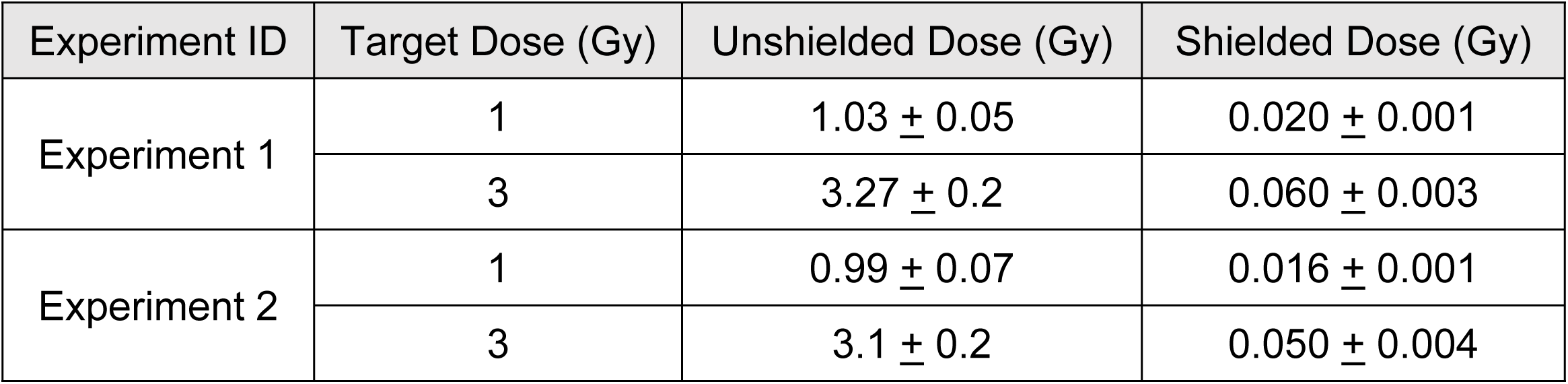
Target and Measured Radiation Doses for Replicate Phenomics Experiments.

### Image-based morphological profiling and radiation science

For this effort, traditional IBMP approaches have been adapted to study the effects of radiation on physiologically relevant cell types. An experimental protocol was developed to irradiate human-derived fibroblasts at target doses of bremsstrahlung radiation in a 96-well format (**Figure 1A**). Following radiation, cells were incubated for varying lengths of time to capture the temporal effects of the radiation damage and recovery cycle. At the desired time point, cells were fixed, stained, and imaged in high-throughput, in accordance with published Cell Painting protocols^6,15^ (**Figure 1B**). The resulting high-content images (HCIs) were segmented down to the single-cell level and vectorized phenoprints were extracted for individual cells. Cell-level phenoprints were processed and averaged to generate aggregate well-level morphological profiles. These mean well-level phenoprints were then used to compare across radiation conditions.

**Figure 1.**
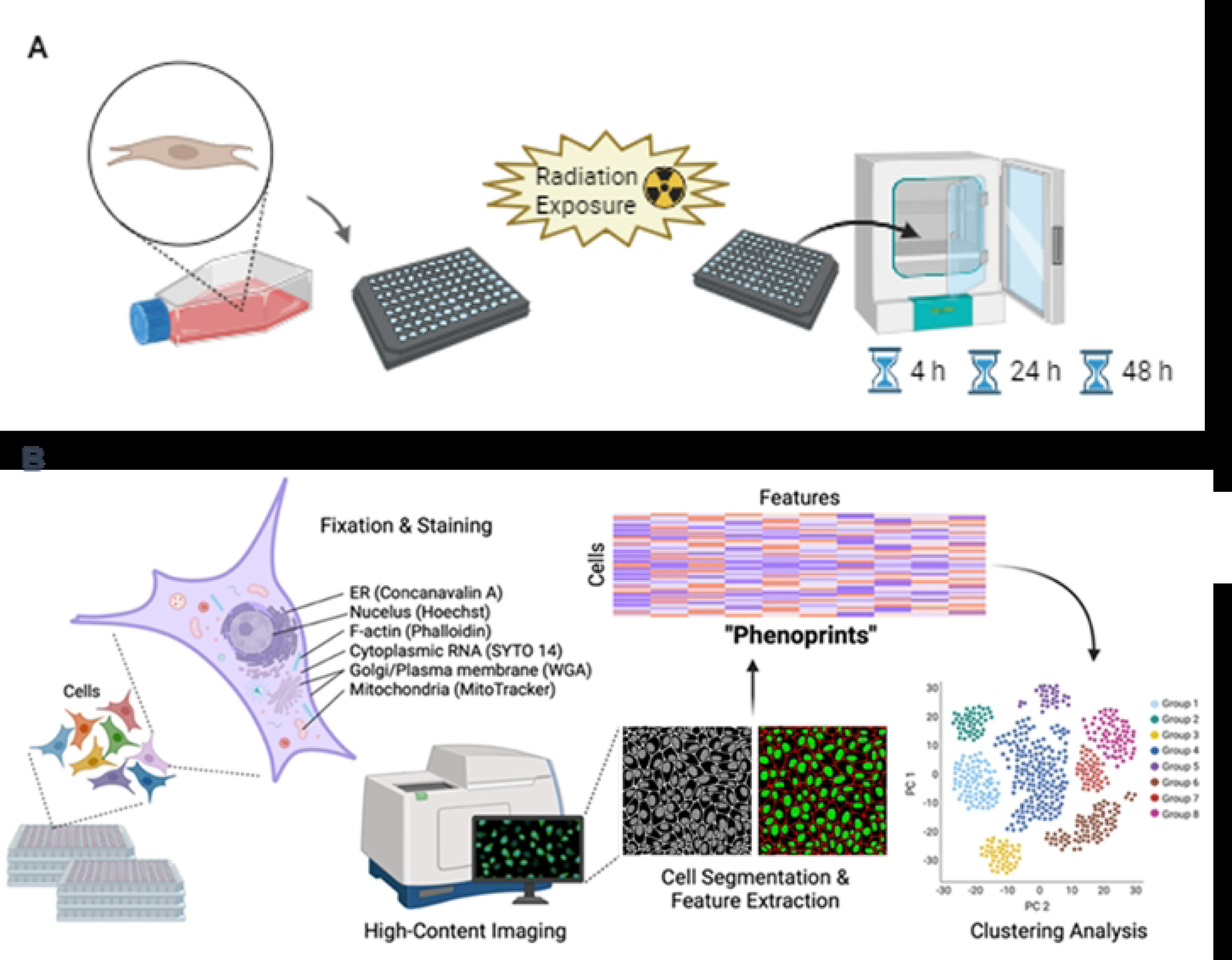
Cell Painting-Based Morphological Profiling for Radiation-Exposed Cells. **A**. Fibroblast cells were cultured in 96-well microplate format prior to being exposed to radiation. After exposure, the cells were incubated for a fixed period of time prior to undergoing Cell Painting. **B**. Exposed and control cells were fixed, permeabilized, and stained using a set of six Cell Painting dyes. The cells were then imaged in high-throughput using a confocal microscope. The resulting images were segmented to extract phenoprints (i.e., a phenotypic morphological profile) of healthy and radiation exposed cells. Phenoprints were aggregated at the well-level and unsupervised clustering was performed to determine morphological similarities and differences between experimental conditions. Figure adapted from reference^5^. Created in BioRender.

### IBMP captures temporal dose-dependent morphological changes in irradiated fibroblasts

In order to capture the temporal- and dose-based effects of radiation, irradiated fibroblasts were incubated for 4, 24, or 48 hours prior to staining and imaging. Confocal imaging revealed increasing cell density over time in both shielded and radiation-exposed conditions (**Figure 2A**) suggesting that the tested doses of radiation were not high enough to completely kill the cells in exposed samples. Despite slight experimental variation in initial cell counts, in both replicate experiments, cell populations increased at a faster rate in shielded conditions, with dose-dependent growth deficits observed in fibroblasts exposed with 1 Gy and 3 Gy of bremsstrahlung radiation at the 24-hour timepoint. (**Figure 2B**, **Supplemental Dataset S1**). There were no observed statistically significant differences in cell counts at the 4-hour timepoint for either dose. It should be noted that the reported cell counts from shielded images taken 48 hours post-exposure in Experiment 1 are likely higher than reported, as a large number of these images were too dense to reliably perform cell segmentation (**Error! Reference source not found.**). Cumulatively, these results suggest that the selected radiation doses were sufficient to reliably impact cell populations across two experiments in a dose-dependent manner.

**Figure 2.**
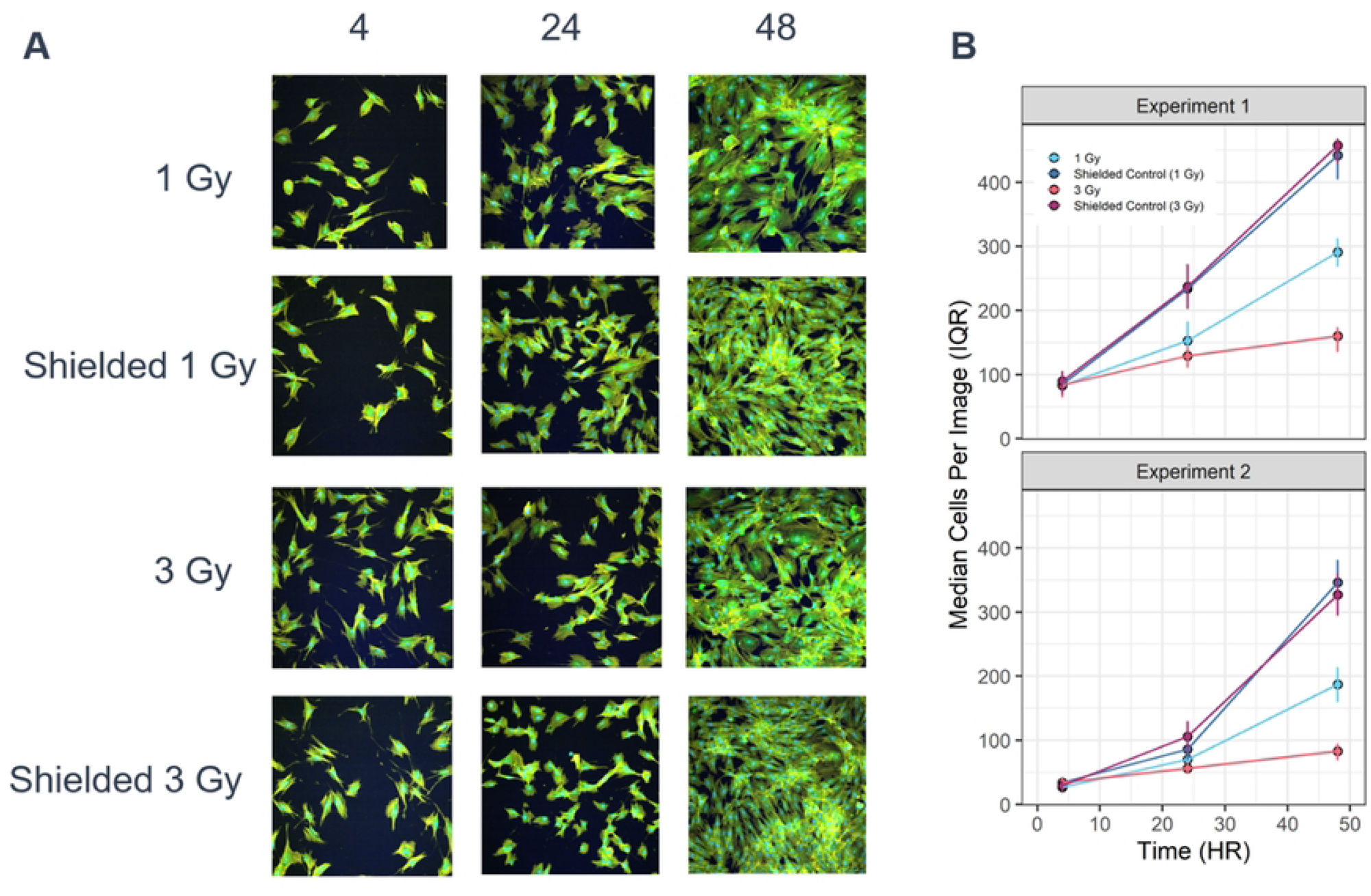
Time- and Dose-Dependent Impacts of Radiation Exposure on Primary Fibroblasts. **A.** Representative Cell Painting images of fibroblasts exposed to, or shielded from, two doses of radiation. Samples were incubated for 4, 24, or 48 hours. Three of the six imaged channels are shown (blue = DNA, green = RNA, yellow = Actin). Images are from Experiment 2 **B.** Median number of segmented cells per image post-QC. Error bars denote interquartile range (IQR).

Beyond cell counts, aggregated well-level morphological profiles, or “phenoprints” were extracted from the two replicate experiments to determine whether radiation elicited distinguishable changes in cell morphology (**Error! Reference source not found.**). The extracted phenoprints consisted of 1,352 measurements per cell and described various properties of cell shape, as well as information about stained subcellular structures (**Supplemental Dataset S2**). To compare phenoprints across conditions, the Pairwise Euclidean distances of mean-centered and variance-scaled phenoprints derived from exposed and unexposed (shielded) wells were calculated. When compared to phenoprint distances within unexposed wells, unexposed vs. exposed distances were significantly different across all conditions (adjusted *P-*value < 0.05), indicating that even after four hours, when cell counts were relatively unchanged, exposed cells had already begun to show the effects of radiation exposure (**Figure 3A, Supplemental Dataset S3**). Exposed and unexposed phenoprints continued to diverge at 24 hours across all dosages. At 48 hours, an interesting phenomenon occurs in which exposed phenoprints appear to recover and better resemble unexposed conditions. This recovery was only consistently observed at the 1 Gy condition. These patterns were also shown using Principal Component Analysis (PCA) clustering (**Figure 3B**). Overall, it appears that unsupervised comparisons of high-dimensional phenoprints are sufficient to capture time and dose-dependent shifts in cell morphology in irradiated cells, even capturing components of cellular recovery.

**Figure 3.**
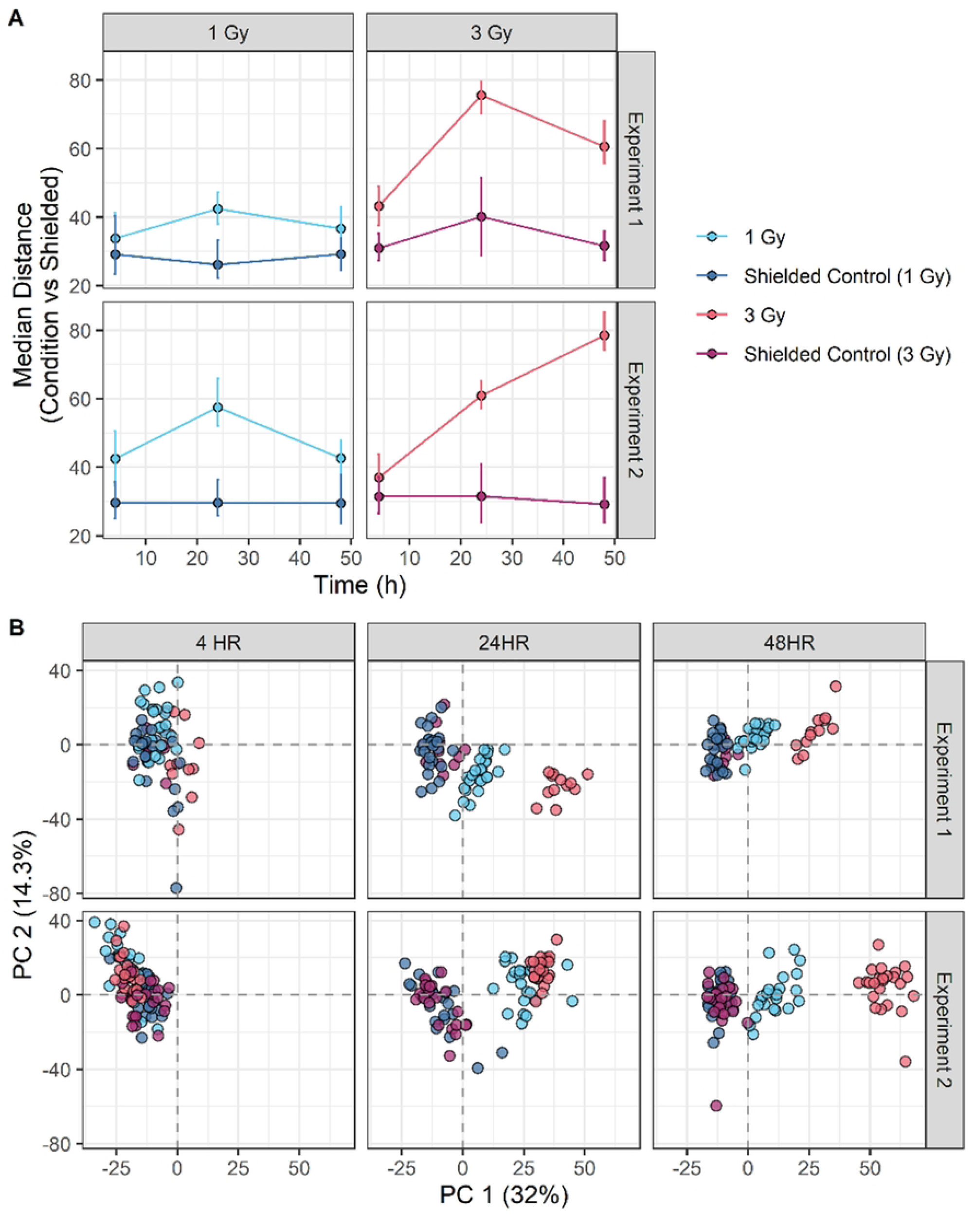
Comparison of High-Dimensional Well-Level Phenoprints across Radiation Exposure Conditions. **A.** Median pairwise Euclidean distances between shielded wells and wells of each exposure condition. Phenoprints were mean-centered and variance-scaled prior to distance calculations. Each condition was compared to shielded wells from the same plate. Shielded wells from the same plate were compared to one another. Error bars denote inner quartile range. All exposed vs shielded distributions are significantly different from shielded vs shielded conditions within a given subplot and time point (Wilcoxon Test; Benjamini-Hochberg Corrected *P-*Value < 0.05) **B.** Principal component analysis (PCA) clustering of normalized well-level phenoprints. Data was mean-centered and variance-scaled prior to clustering. Percent variance along principal component 1 (PC1) and principal component 2 (PC2) are shown. Clustering was performed across all time points and experiments and has been sub-plotted for visualization purposes.

### Morphological profiles reveal temporal impacts of radiation on cellular structure

Despite the fact that morphological profiles consist of a vast number of abstract measurements that represent the whole cell, it was still possible to glean interpretable information about which aspects of cellular morphology might be driving observed changes between conditions. To better understand the morphological drivers of differences observed in the temporal and dose-dependent radiation experiments, feature importance scores were extracted from ensembles of random forest models trained to classify irradiated cells in six independent conditions (**Table 2**). In this analysis, phenoprint features with higher importance scores more strongly impact the performance of a random forest model. While subject to bias from high cardinality variables, importance scores can provide a starting point for hypothesis generation and can be used to down-select conditions for more targeted molecular analyses.

**Table 2.**
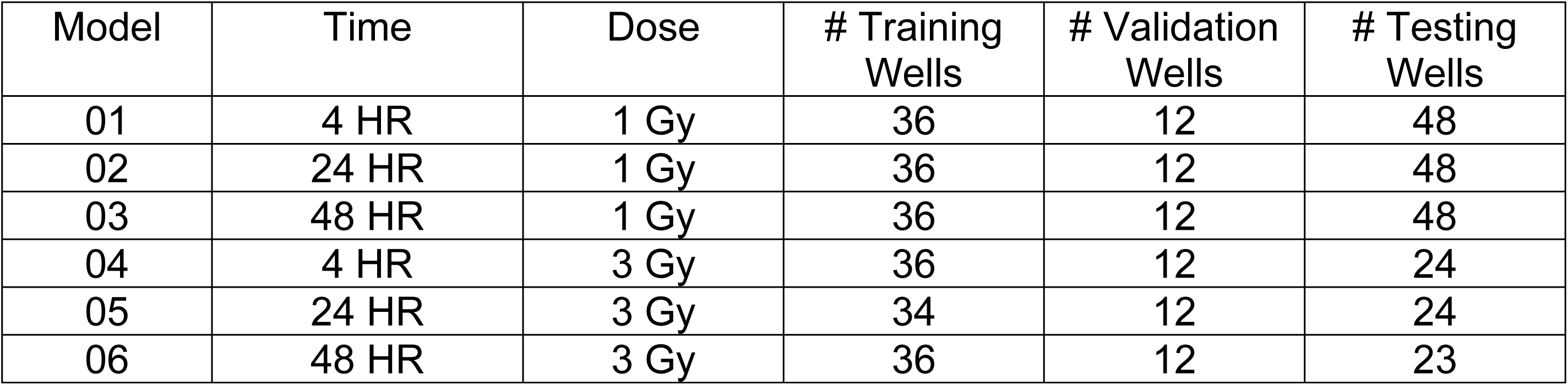
Summary of Random Forest Training, Validation, and Testing Data. Models were trained and validated on a 75:25 split of Experiment 2 data and tested on all data from Experiment 1. For each model ID, an ensemble of 100 random forest models were trained. The same data splits were used across a given model ensemble. All models were trained to classify irradiated from control wells.

| Model | Time | Dose | # Training Wells | # Validation Wells | # Testing Wells |
| --- | --- | --- | --- | --- | --- |
| 01 | 4 HR | 1 Gy | 36 | 12 | 48 |
| 02 | 24 HR | 1 Gy | 36 | 12 | 48 |
| 03 | 48 HR | 1 Gy | 36 | 12 | 48 |
| 04 | 4 HR | 3 Gy | 36 | 12 | 24 |
| 05 | 24 HR | 3 Gy | 34 | 12 | 24 |
| 06 | 48 HR | 3 Gy | 36 | 12 | 23 |

For a given random forest model, the importance scores of features related to each imaged structure (*e.g.,* DNA, RNA, Actin, ER, Golgi, Mitochondria) as well as measurements related to shape (*e.g.,* AreaShape) and cell density (*e.g.,* Neighbors) were summed to understand the cumulative impact of each feature type on the ability to identify profiles of irradiated cells. For each model condition, cumulative importance scores were averaged across 100 models. The importance of each feature type was found to be generally consistent across radiation dose, with feature types varying based on time post-exposure (**Figure 4A**). The importance of cell area and number of neighbors increased over time, indicating that profiles are at least to some degree impacted by the effect of radiation on cell growth and subsequent density. However, importance was observed in other features, indicating that the profiles were not solely driven by growth related changes. For example, the importance of features related to the endoplasmic reticulum (ER) peaked at 24 hours post-exposure, suggesting that the ER may have been morphologically impacted by radiation at this time point, prior to partially recovering by 48 hours post-exposure. Additional experimentation is required to determine whether the decrease in the importance of ER-related features at 48 hours post-exposure is due to the return of the ER to a control-like state or whether the importance of ER-related features is instead just outweighed by other more substantial changes found at 48 hours.

**Figure 4.**
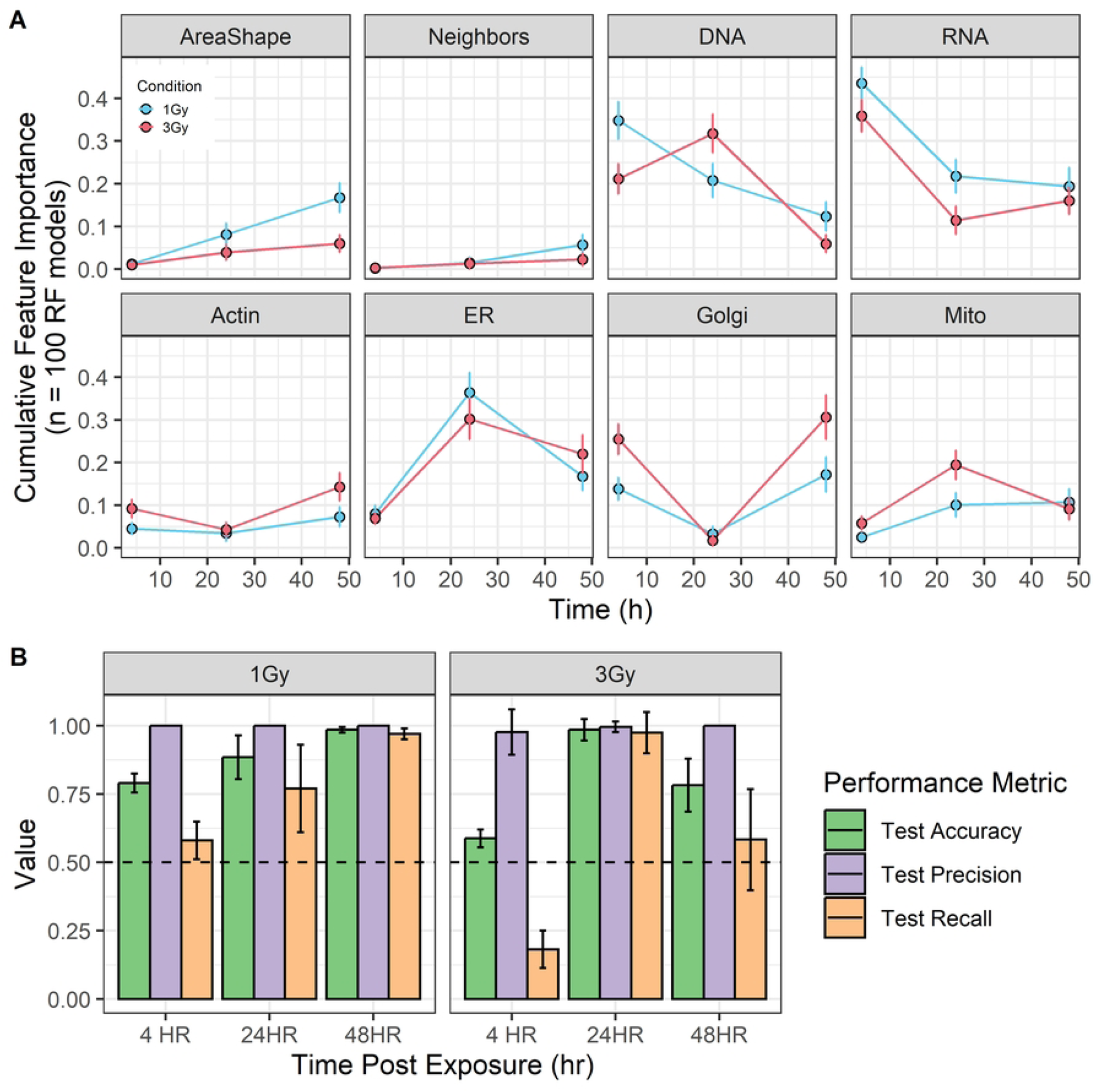
Random Forest Feature Importance and Performance. Ensembles of 100 random forest models trained to classify irradiated from control well-level phenoprints in six conditions (*e.g.,* 1 Gy 4 hours, 1 Gy 24 hours, 1 Gy 48 hours, 3 Gy 4 hours, 3 Gy 24 hours, 3 Gy 48 hours). **A)** Cumulative feature importance across eight feature categories (*e.g.,* six channels, AreaShape, Neighbors). Feature importance was summed within each individual model. For each condition, the mean and standard deviation of importance is reported across 100 models. Some features contain information on multiple channels and therefore it is possible for the total cumulative importance to be greater than one for a given condition. **B)** Performance of each model on test data. Mean and standard deviation of three performance metrics taken across 100-model ensembles are reported for each condition. Dashed line denotes a random performance of 0.5.

As a final analysis, ensembles of random forest models were used to evaluate experimental variability across exposure doses and post-exposure time points. Specifically, the ability of models trained on Experiment 2 to identify irradiated morphological profiles from Experiment 1 was tested. Models generally performed well on test data from Experiment 1, especially at the 24-hour time point (**Figure 4B**). Unsurprisingly, models were the least generalizable at the four-hour time point, where phenoprint signatures were relatively weak and noisy (**Figure 3B**). Model recall of the 48 hour, 3 Gy condition was also relatively poor, which is likely a result of training on Experiment 2, which had a more robust phenomic signature at this time point than in Experiment 1, indicating some experimental variability. Overall, IBMP in combination with random forest or other interpretable machine learning approaches can be used to help identify radiation conditions that elicit reproducible effects and predict subcellular changes impacting these effects.

## DISCUSSION

Recently, phenomics-based analytical approaches have shown utility in accelerating drug discovery pipelines and advancing various fields of basic research^4,14,16^. Here, the utility of applying phenomics-based approaches to studying the effects of ionizing radiation exposures is demonstrated. Experimentally, a procedure was optimized to use an in-house linear accelerator to deliver controlled doses of bremsstrahlung radiation evenly across 96-well cell culture plates. Human-derived fibroblasts were exposed to two different doses of radiation and monitored at 4-, 24-, and 48-hours post irradiation. The irradiated cells were stained for six cellular structures (DNA, RNA, Golgi, ER, actin, and mitochondria) and imaged in high-throughput. High-dimensional quantitative morphological profiles, or “phenoprints,” were generated at the cell-level and aggregated to the well-level. Unsupervised clustering of well-level phenoprints revealed time- and dose-dependent changes in the morphology of irradiated cells relative to shielded controls (**Figure 3B**). Phenomic differences between unexposed and irradiated cells could be observed as early as 4 hours post-exposure, prior to any difference in cell counts resulting from exposure, though the differences grew stronger over time. Furthermore, radiation effects could be captured by morphological changes in different cellular structures over time (**Figure 4A**), with the ability to observe cellular rebounding to the unexposed state.

Overall, these results demonstrate the utility of Cell Painting and IBMP to identify radiation conditions which impact cell morphology. This pipeline could serve as a rapid pre-screening approach to down-select conditions for follow-up with low-throughput mechanistic assays. This would ensure more complex experimentation were conducted at exposures and time points that were the most physiologically relevant. Additionally, this work suggests that Cell Painting and IBMP could be used as a method to screen for radiation exposure prophylactics or therapeutic treatments. Chemical countermeasures of interest could be administered to the cells pre- or post-exposure to determine if a phenoprint shift could be prevented, or if the cells return to the “healthy” unexposed condition more rapidly post-exposure. The strategy could also be used to assess the effects of repeated exposure and cell resilience. In this work, it was observed that cells exposed to 1 Gy of radiation had started to resemble unexposed controls by 48 hours, while recovery following a 3 Gy exposure was inconsistent across experimental replicates. This demonstrated that the ability to rebound from a single insult is dose-dependent. The impact of multiple exposures could be explored by exposing cells, allowing them to recover, and then exposing them again to the same condition and observing whether the same recovery is observed. Such experimentation could be an efficient way to gain a deeper insight into chronic radiation exposure.

One limitation of this work is that there was not a clear approach to control for the impacts of cell density on well-level phenoprints. Shielded controls cells exhibited higher cell counts (growth rates) than irradiated cells at 24- and 48-hour time points (**Figure 2B**), and while this change in cell density was likely a direct impact of radiation treatment, the extent to which cell density impacted observed differences in irradiated samples and the subsequent phenoprint recovery at 48 hours with a 1 Gy exposure remains to be determined.

Overall, this work demonstrates the utility of morphological profiling in radiation science. Beyond using Cell Painting and IBMP to characterize the morphological impacts of radiation exposure, the outlined approach has the potential to be applied to down-select conditions for low-throughput mechanistic assays as well as to address key questions related to repeated radiation exposures and mitigation options. Importantly, target doses sufficient to elicit reproducible morphological changes in human-derived fibroblasts were identified. Lower exposure doses, singular or repeated, may still be able to elicit responses and should be investigated further. Future work should aim to combine these conditions with the power of high-throughput IBMP drug-screening based approaches^4^ to discovery novel countermeasures against the effects of radiation exposure.

## METHODS

### Cell culture

Primary Dermal Fibroblasts (HDFn, PCS-201-010), Fibroblast Basal Medium, Fibroblast Growth Kit, Trypsin-EDTA for Primary Cells, Trypsin Neutralizing Solution, and Dulbecco’s Phosphate Buffered Saline were purchased from ATCC (Manassas, VA). T75 culture flasks (229341) were purchased from CELLTREAT Scientific Products, Hank’s Balanced Salt Solution (HBSS, 14175103) and conical tubes were purchased from Thermo Fisher Scientific. Sterile 96-well Half Area High Content Imaging Film Bottom Microplates (4680) were purchased from Corning (Kennebunk, ME). The fluorescent labeling reagents and used in Cell Painting are described in Table 3.

**Table 3:** Cell Painting Dyes Information.

| **Organelle** | **Label** | **Excitation/Emission**<br>**Wavelength (nm)** | **Vendor** | **Cat-No** |
| --- | --- | --- | --- | --- |
| Mitochondria | Mitrotracker | 644/665 | Thermo Fisher Scientific | M22426 |
| Nucleus | Hoechst 33342 | 350/461 | Thermo Fisher Scientific | H1399 |
| Nucleoli/RNA | SYTO 14 Green | 517-521/547-549 | Thermo Fisher Scientific | S7576 |
| ER | Concanavalin A<br>AlexaFluorTM 488 | 495/519 | Thermo Fisher Scientific | C11252 |
| Actin | Phalloidin<br>AlexaFluorTM 568 | 578/600 | Thermo Fisher Scientific | A12380 |
| Plasma<br>Membrane | Wheat Germ<br>Agglutinin<br>AlexaFluorTM 555<br>Conjugate | 555/565 | Thermo Fisher Scientific | W32464 |

Complete Fibroblast Media was prepared by adding the contents of the Fibroblast Growth Kit to the Fibroblast Basal Medium and this complete media was subsequently used in all culture steps. Upon receipt, HDFn cells were thawed and cultured at 37°C and 5% CO_2_. After the HDFn cells reached 80% confluence, cells were rinsed with PBS and removed from the flask with trypsin. HDFn cells were counted and stocks of 500,000 HDFn cells per mL of preservation media (80% Complete Fibroblast Media, 10% DMSO, 10% FBS) were created and stored in cryovials at -180°C.

Each experimental replicate required one HDFn cell vial to be thawed, cultured to Day 4 in a T75 flask, then expanded through one consecutive passage to achieve a cell density that was sufficient to seed 1,000 cells per well of a 96-well plate. Edge wells of the plate were seeded but not used in Cell Painting or analysis to eliminate any potential edge effects. Seeded plates were centrifuged at 500xg for 1 minutes then incubated overnight until radiation exposure on the following day.

### Radiation treatment

Cell plates were transferred from the incubator to a thermally-insulated container until radiation experiments were completed. An irradiating bremsstrahlung beam was produced by impinging 12 MeV electrons on to a 2 mm thick tungsten radiator mounted on a 2.5 cm thick water-cooled aluminum block, absorbing primary electrons that traverse the tungsten (**Figure 5A-B**). The microplates with the fibroblasts were placed 2.5 m from the radiator ensuring a uniform dose profile over a 6 cm diameter. To ensure secondary electron equilibrium, a 9.5 mm thick polyethylene plate was directly in front of the microplates. Half of the microplate and polyethylene plate were shielded by 20.3 cm of lead. A Monte Carlo simulation was conducted to determine the thickness of lead required to block the majority of the radiation dose. In order to limit edge effects, border wells and wells at the edge of the lead shielding were seeded, but were omitted from imaging and downstream analysis (**Figure 5C**). The fibroblasts on the unshielded side received the full radiation dose, while the fibroblasts on the shielded side received approximately 1-2% of the full dose, as seen in **Figure 5D**. The unshielded dose rate was monitored in real-time by a 0.6 cm^3^ ionization chamber (RadCal 10X6-0.6 ionization chamber connected to an ADDM+ digitizer). The unshielded and shielded total delivered doses were measured by a series of optically stimulated luminescence dosimeters placed on the cell plate behind the 9.5 mm polyethylene sheet for electron equilibrium. After irradiation, cell plates were incubated at 37°C for 4, 24, and 48 hours prior to analysis.

**Figure 5:**
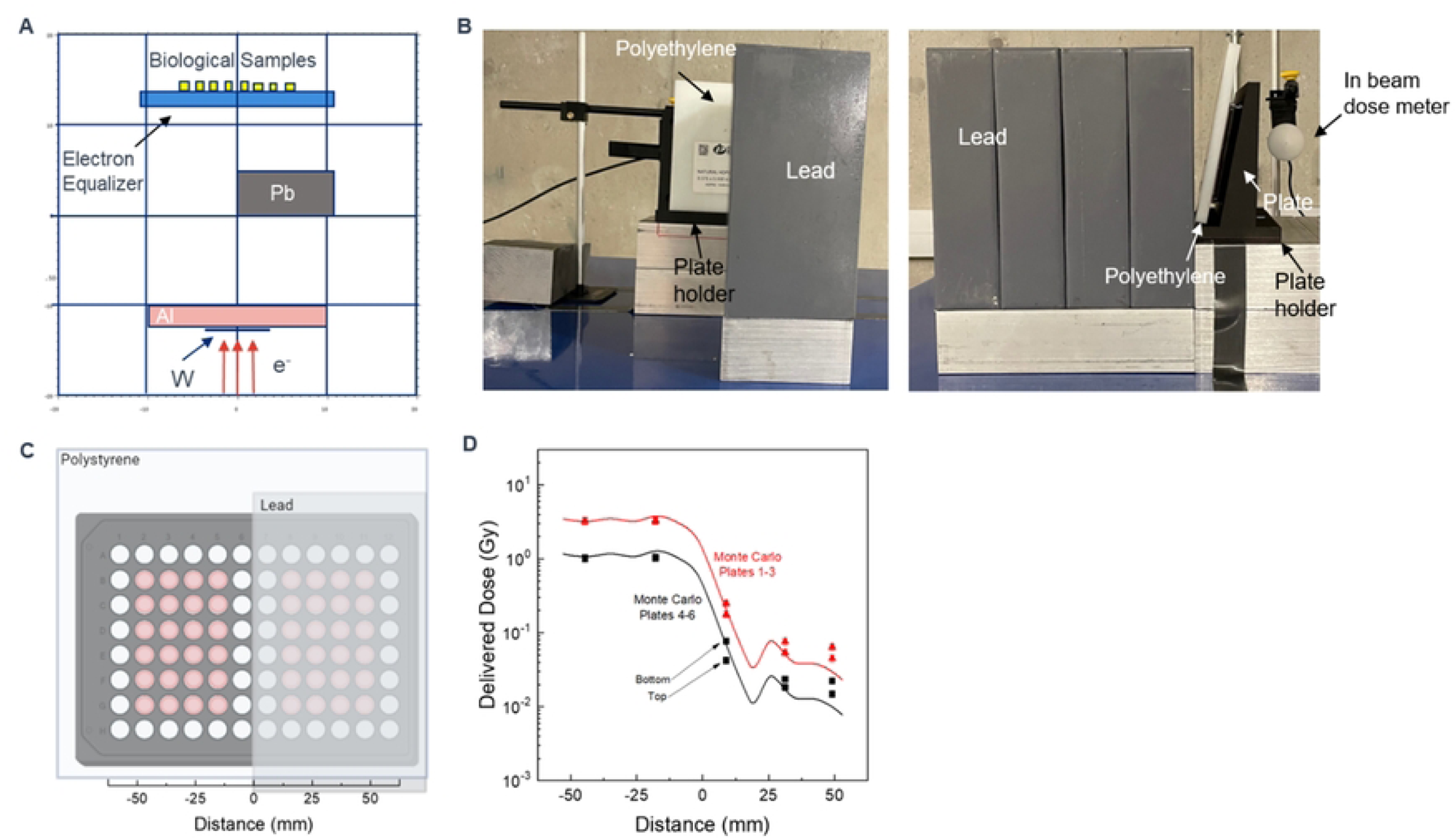
Radiation Exposure Setup. **A**. Overhead schematic of radiation exposure setup used for experimental exposures, including an electron source, tungsten radiator, water cooled aluminum (Al) block, lead shielding (Pb), and polyethylene electron equalizer. **B**. Representative photos of the experimental setup from the front (left) and side (right) view. **C**. Plate design for radiation experiments. To minimize plate effects, edge wells and wells at the shielding interface were omitted from imaging and downstream processing (white = not imaged, pink = imaged). Distance in millimeters from the center of the plate was provided for comparison to simulation results. Created in BioRender. **D**. Simulated (lines) vs measured (points) radiation doses across 96-well plates targeted to receive 1 Gy (black) or 3 Gy (red) of radiation. Measured doses were taken from Experiment 1. Top and bottom designations denote radiation exposure levels at the top and bottom of three stacked plates. Plates were shielded at distances greater than 0 mm.

### Cell painting and image acquisition

At 4-, 24-, and 48-hours post-irradiation, the microplates were removed from the incubator and prepared for Cell Painting. MitoTracker dye was diluted in Complete Fibroblast Media at a 1:2000 dilution factor. Media from the cell plates was removed and 100 µL of diluted MitoTracker was added to the wells and incubated at 37°C for 30 minutes. Following incubation, 25 µL of 16% paraformaldehyde was added to each well and the cell plate was incubated for 20 minutes at room temperature in the dark. The plate was washed in triplicate with HBSS and 100 µL of 0.1% Triton-X was added to each well. After incubation for 15 minutes at room temperature in the dark, the plate was washed in triplicate with HBSS.

Cell Painting was performed, as previously described^6^. Briefly, a Cell Painting dye master mix consisting of Hoechst 33342, SYTO 14 Green, Concanavalin A AlexaFluor^TM^ 488, Phalloidin AlexaFluor^TM^ 568 and Wheat Germ Agglutinin AlexaFluor^TM^ 555 Conjugate was made in a HBSS + 1% bovine serum albumin (w/v) solution. Cells were stained with 50 µL of the master mix for thirty minutes at room temperature after being centrifuged at 500 x *g* for 1 minute to remove bubbles. Following staining, the cells were washed three times with 100 µL of HBSS and covered in foil. The plates were imaged using an ImageXpress Micro Confocal High-Content Imaging System (Molecular Devices) in confocal mode (60μm slit) with a 20X objective. Twelve sites per well were acquired with 6 channels per site. The following bandpass filters were used to visualize the channels: FF01-452/30, FF01-562/40, FF02-520/28, FF01-624/40, FF01-595/31 and FF01-681/24.

### Cell segmentation and generation of phenoprints

Following Cell Painting and raw image acquisition, images were processed and segmented using CellProfiler v4.2.1^8^. CellProfiler is a free bio imaging software and was selected for use due to its wide variety of features and ease of optimization. A public CellProfiler pipeline was adapted for in-house images^6^. Three scripts were run in sequence to perform 1) image-level quality control measurements (QC), 2) illumination correction, and 3) cell segmentation.

In brief, images from each channel were illumination-corrected to address differences in image illumination (i.e., fluorescence intensity), which occurs naturally due to microscope optics. Cells from each image were then segmented by first identifying the nucleus from the DNA channel and then finding the cytoplasm boundaries from the RNA channel. For each cell, >1200 morphological measurements were collected, cumulatively referred to as single-cell phenoprints. The final outputs of the CellProfiler pipeline were 1) quantitative single-cell phenoprints for all images, 2) images overlaid with outlined cell segmentations used for manual QC, and 3) QC measurements for each segmented image.

### Phenoprint quality control, normalization, and aggregation

Immediately following segmentation, mis-segmented images were manually identified and flagged for removal. An in-house phenomics pipeline was then applied to process cell-level phenoprints. Cell-level phenoprints were annotated with image metadata (*e.g.,* radiation conditions, time points).

Following removal of mis-segmented images, phenoprints underwent additional quality control. Cells in the top or bottom 1% of measured cell areas were removed in an effort to exclude mis-segmented cells in otherwise well-segmented images. Cells coming from wells with fewer than 10 cells were additionally removed as such a small number of cells cannot be reliably summarized into a well-level phenoprint. Following quality control, cell counts per image were calculated (**Supplementary Dataset S1**) and well-level phenoprints were generated by calculating the average phenoprint of all cells in each well.

Finally, well-level phenoprints were normalized by the negative controls on each plate^17^. In brief, for each plate, the median and median absolute deviation (MAD) of each morphological measurement across the negative controls (*e.g.,* shielded cells) were calculated. MAD normalization was applied to all data within each plate by subtracting the median and dividing by the MAD:

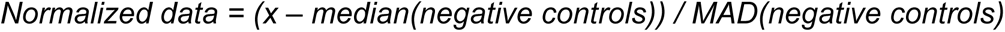

As a result of this normalization, each measurement in the phenoprints of shielded cells within a given plate had a median of zero and a MAD of one. This allows a fairer comparison of the impact of radiation across multiple experiments. Processed well-level phenoprints were used in downstream phenomics analyses (**Supplementary Dataset S2**).

### Distance measurements and unsupervised clustering of well-level phenoprints

Euclidean distances of mean-centered and variance scaled well-level phenoprints were measured between all pairs of shielded wells and all pairs of shielded and irradiated wells within a single experiment, exposure (*e.g.,* 1 Gy, 3 Gy), and time point. Statistical comparisons between shielded-shielded and shielded-irradiated distance distributions were made with the nonparametric Wilcoxon rank sum test. P-values were adjusted for multiple corrections with the Benjamini-Hochberg procedure^18^. Principal component analysis (PCA) was performed on mean-centered and variance-scaled well-level phenoprints with the built in “princomp” function in R version 4.1.3. Unless otherwise stated, variables that were non-numeric, contained numeric metadata, contained NA values, or had zero variance were removed prior to clustering. Additionally, only variables shared across phenoprints in replicate experiments were retained to allow for a more representative comparison between the two datasets. The percent of variance captured along the first two principal components was reported.

### Random forest models and cumulative feature importance

Data splits and random forest models were implemented with the scikit-learn package in Python^19^. Because it was a smaller dataset, Experiment 1 was set aside as a final test set and was excluded from model training and validation. Well-level phenoprints from individual time points and radiation doses of Experiment 2 were split in to training and validation sets at a 75:25 ratio, stratified to maintain consistent ratios of class labels (*e.g.,* irradiated vs control).

Random forest models were trained to differentiate between irradiated and control well-level phenoprints for each time point and radiation dose, resulting in a total of six conditions (**Table 2**). Random forest was implemented in scikit-learn with the default settings (*e.g.,* 100 trees). To address potential variability that could occur given the small training set size, 100 random forest models (*e.g.,* unique seeds) were generated for each of the six conditions. Feature importance scores were recorded for each of the 600 models (**Supplementary Dataset S4**). To summarize the cumulative importance of a cellular structure or specific feature type, feature importance scores were summed across each channel (*e.g.,* DNA, RNA, actin, Golgi, ER, mitochondria) as well as across features having to do with Area Shape or number of neighbors. The mean and standard deviation of cumulative importance were reported for each of the six classification conditions. Correlation measurements relate two cellular structures and therefore the importance of each correlation feature was counted toward both structures. This resulted in a cumulative importance greater than one.

Finally, for each condition, the performance of the 100 random forest models were evaluated on test data sets for that condition. The test data consisted of well-level phenoprints from an independent experiment (Experiment 1). Prediction accuracy, precision, and recall were determined with scikit-learn (**Supplementary Dataset S5**). Mean and standard deviation of these metrics were calculated across each 100-model ensemble. All random forest results were visualized in R.

## Data availability

Processed well-level phenoprints, final cell counts per image, statistical comparisons, and random forest feature information and performance metrics are available in the supplemental materials (**Supplementary Datasets S1-S5**).

## ACKNOWLEDGMENTS

This work was supported by internal funding provided by the Johns Hopkins University Applied Physics Laboratory (to O.N.T.).

## AUTHOR CONTRIBUTIONS

Cell culture, Cell Painting staining, and image acquisition was performed by O.N.T and S.M.T. A.W.H. was responsible for all radiation modeling and exposures. L.T performed image segmentation using CellProfiler and L.J.D devised and implemented phenomics analyses. R.J. M contributed to the manuscript and experimental protocol development. O.N.T together with S.M.T, A.W.H, and L.J.D designed the experiments, interpreted the data, and wrote the manuscript.

## COMPETING FINANCIAL INTERESTS

All authors declare no competing financial interests.

## Notes

### Competing Interest Statement

The authors have declared no competing interest.

